# Habitual Stone-Tool Aided Extractive Foraging in White-Faced Capuchins, *Cebus Capucinus*

**DOI:** 10.1101/351619

**Authors:** Brendan J Barrett, Claudio M Monteza-moreno, Tamara DOGANDŽIĆ, Nicolas Zwyns, Alicia IBÁÑEZ, Margaret C Crofoot

**Affiliations:** Cognitive and Cultural Ecology Group, Max Planck Institute for Ornithology, Radolfzell, Germany; Department of Human Behavior, Ecology, and Culture, Max Planck Institute for Evolutionary Anthropology, Leipzig, Germany; Animal Behavior Graduate Group, University of California, Davis. Davis, CA. USA; Department of Anthropology, University of California, Davis. Davis, CA. USA; Smithsonian Tropical Research Institute, Balboa, Ancón, Panamá, Republic of Panamá; Department of Human Evolution, Max Planck Institute for Evolutionary Anthropology, Leipzig, Germany; Department of Anthropology, University of Pennsylvania, Philadelphia, PA. USA; Estación Científica Coiba AIP, Ciudad del Saber, Panamá, Republic of Panamá; Jadwin Av. 108B, Gamboa. Republic of Panamá

## Abstract

Habitual reliance on tool use is a marked behavioral difference between wild robust (genus *Sapajus*) and gracile (genus *Cebus*) capuchin monkeys. Despite being well studied and having a rich repertoire of social and extractive foraging traditions, *Cebus sp* have rarely been observed engaging in tool use and have never been reported to use stone tools. In contrast, habitual tool use and stone-tool use by *Sapajus* is widespread. We discuss factors which might explain these differences in patterns of tool use between *Cebus* and *Sapajus*. We then report the first case of habitual stone-tool use in a gracile capuchin: a population of white-faced capuchins (*Cebus capucinus imitator*) in Coiba National Park, Panama who habitually rely on hammerstone and anvil tool use to access structurally protected food items in coastal areas including *Terminalia catappa* seeds, hermit crabs, marine snails, terrestrial crabs, and other items. This behavior has persisted on one island in Coiba National Park since at least 2004. From one year of camera trapping, we found that stone tool use is strongly male-biased. Of the 205 unique camera-trap-days where tool use was recorded, adult females were never observed to use stone-tools, although they were frequently recorded at the sites and engaged in scrounging behavior. Stone-tool use occurs year-round in this population, and over half of all identifiable individuals were observed participating. At the most active tool use site, 83.2% of days where capuchins were sighted corresponded with tool use. Capuchins inhabiting the Coiba archipelago are highly terrestrial, under decreased predation pressure and potentially experience resource limitation compared to mainland populations– three conditions considered important for the evolution of stone tool use. White-faced capuchin tool use in Coiba National Park thus offers unique opportunities to explore the ecological drivers and evolutionary underpinnings of stone tool use in a comparative within- and between-species context.

## 1. INTRODUCTION

Extractive foraging permits many generalist species to access structurally protected resources. It requires both physiological specializations and/or cognitive traits that aid in resource manipulation and extraction. Extractive foraging may permit organisms to outcompete species which lack such adaptations and may be of great importance in ecologically stressful periods as a fallback foraging strategy (Marshall and Wrangham, 2007). It is also important to the ecological success and evolutionary history of many primates.

Tool use is a taxonomically widespread (Bentley-Condit and Smith, 2010; Shumaker et al., 2011) form of extractive foraging which may permit access to novel resources, expand diet breadth, and may be useful to more efficiently or safely access existing resources. Comparative studies of tool use are key to understanding the ecological and social factors that drive its evolution within the primate lineage. In primates, the cognitive skills required for tool use was likely selected for alongside social and ecological intelligence (Reader et al., 2011) and generalized problem solving (Seed and Byrne, 2010) (but see Teschke et al. (2013) for exceptions to this in other taxa). Once tool use evolves it may create interesting eco-evolutionary feedbacks via gene-culture coevolution and/or cultural niche construction that are likely important in hominin evolution (Richerson and Boyd, 2005; Kacelnik, 2009; Laland et al., 2010). The cultural transmission of toolkits and associated behaviors is of central importance to human evolution; it is key to our success as the most-widely dispersed vertebrate on Earth.

Comparative studies of stone tool use in non-human primates is of particular interest to paleoanthropologists as it provides a model of the possible tool use behaviors that led to the emergence of earliest known stone tool production in hominins. Percussive techniques like those observed in non-human primates may have been the precursor to the earliest hominin stone tool making around 3 mya (Panger et al., 2002a; Toth and Schick, 2009; McPherron et al., 2010; Harmand et al., 2015; Arroyo and de la Torre, 2016). Thus, a better understanding of tool use in non-human primates helps us to validate and critically evaluate our interpretations of the fossil and archaeological records (Proffitt et al., 2016). Further, comparative studies of tool use in extant primates allow us to better interpret and understand what early lithic technologies looked like, how they may have been used by ancient hominins, as well as how site preservation and visibility (e.g. Panger et al. (2002a)) affect our ability to interpret the fossil record.

### 1.1. Differences in tool use between robust and gracile capuchins

Among capuchin monkeys, habitual reliance on tools, and stone tools in particular, has been considered a distinguishing feature of the larger-bodied robust capuchins (*Sapajus*) from the smaller gracile capuchins (*Cebus*) since their divergence 6.5 mya (Lynch Alfaro et al., 2012). Stone tool use has been observed in the wild in all well-studied robust capuchins species including black-striped capuchins, *Sapajus libidinosus* (Ottoni and Mannu, 2001; Fragaszy et al., 2004a; Mendes et al., 2015); yellow-breasted capuchins, *Sapajus xanthosternos* (Canale et al., 2009); blonde capuchins, (*Sapajus flaviius*) (Ferreira et al., 2009); black-horned capuchins, *Sapajus nigritus* (Rocha et al., 1998); and black-capped capuchins, *Sapajus apella* (Boinski et al., 2000) (based off of taxonomic reclassifications (Boubli et al., 2012; Lynch Alfaro et al., 2012) and reviews by Ottoni and Izar (2008) and Garber et al. (2012)). Other forms of tool use, especially the use of sticks to dig and probe, have also been reported in *Sapajus* (Fernandes, 1991; Souto et al., 2011; Torralvo et al., 2017).

Using one liberal definition of tool use adapted from Chevalier-Skolnikoff (1989) (“the use of one unattached object to effect a change in another”), many of the combinatorial actions that are habitual components of *Cebus* species’ behavioral repertoire would qualify as tool use (see Fragaszy et al. (2004b) chapter 7 for a review). This includes common foraging behaviors like pounding and scrubbing fruits and animal prey on rocks or branches (Oppenheimer, 1968; Panger et al., 2002b; Perry, 2009; Méndez-Carvajal and Valdés-Díaz, 2017), as well as their use of branches as fulcrums, and breaking sticks in aggressive displays (Chevalier-Skolnikoff, 1990). Although there are several reports of *Cebus* using sticks as clubs, most are one-time observations (Boinski, 1988); whether this behavior is intentional or exploratory is debated (Fragaszy et al., 2004b). This contrasts with Beck’s (1980) more commonly accepted definition where three criteria must be satisfied to qualify as tool use: the potential tool cannot be part of the animal, must not be attached to the surrounding environment, and must be manipulated to achieve a useful outcome (but see Hansell and Ruxton (2008) for inconsistencies regarding use of this definition).

Using this more stringent and widely used definition, tool use is rarely observed in gracile capuchins. Some social groups of white-faced capuchins, *Cebus capucinus imitator*, use leaves as sheathes to process stinging *Automeris* caterpillars and *Sloanea* fruits (Panger et al., 2002b; Perry et al., 2017). Both white-fronted capuchins, *Cebus albifrons*, and white-faced capuchins have been observed to use leaves as sponges for drinking (Phillips, 1998; Perry et al., 2017). However, while tools may be helpful for acquiring these resources, they are not required. Habitual use of tools, and stone tools in particular, is considered one of the defining behavioral differences between *Cebus* and *Sapajus* (Lynch Alfaro et al., 2014). The near absence of tool use in gracile capuchins is puzzling as they are otherwise highly innovative (Perry et al., 2017) and have a wide array of social (Perry et al., 2003) and foraging traditions (Chapman and Fedigan, 1990; Panger et al., 2002b; O’Malley and Fedigan, 2005) which are socially learned (Perry, 2009; Barrett et al., 2017). In addition, many physiological and social traits which are associated with tool use, including tolerance of close observation by conspecifics (Oppenheimer, 1968; Perry and Ordoñez Jimenez, 2006), exploratory behavior (Fragaszy and Visalberghi, 2004; Perry et al., 2017), and dexterous hands (Torigoe, 1985; Costello and Fragaszy, 1988; Christel and Fragaszy, 2000) are present in both gracile and robust capuchins.

### 1.2. Hypotheses explaining the origins and/or maintenance of tool use

Stone tool use has been observed in only three genera of wild primates: chimpanzees (*Pan troglodytes*) (Sugiyama and Koman, 1979; Boesch and Boesch, 1990), robust capuchins (*Sapajus spp.)* (Ottoni and Izar, 2008), and macaques (*Macaca fascicularis*) (Malaivijitnond et al., 2007; Gumert et al., 2009). Within each of these groups, however, significant variation in tool use behavior exists. This variation warrants further explanation.

Fox et al. (1999) synthesized three hypotheses for why primates use tools. The *necessity hypothesis* suggests that ecological need drives stone tool use. The original formulation of this hypotheses presupposes that resources are distributed in a manner that favors scramble competition (which may be driven by high population densities), and predicts that tool use will be common in populations with relatively less favorable energy budgets. The necessity hypothesis primarily pertains to the evolutionary origins of tool use, but could also account for its maintenance. Subsequent formulations of this hypothesis have focused on an overall reduction in resource availability and/or seasonally limited access to food as the causal factor driving tool use innovations (Sanz and Morgan, 2013; Koops et al., 2013). However, current data on environmental seasonality and resource fluctuations may not reflect the historic environmental conditions experienced by species (or population) when tool use traditions emerged. These results must therefore be interpreted with caution.

The *opportunity hypothesis* posits that increased availability of, and encounter rates with, tool material and appropriate food resources drives tool use (Fox et al., 1999). This hypothesis suggests that tool-using populations simply live in conditions where they are more likely to encounter materials for tool use and resources which require tools for extraction. It does not make any assumptions or predictions about differences in resource distributions or foraging efficiency between tool-using and non-tool-using populations.

The *relative profitability hypothesis* was originally articulated in studies of tool use in New Caledonian crows (Rutz et al., 2010; Rutz and St Clair, 2012). It suggests that tool-aided foraging behaviors are used when they have a better energetic return rate for accessing structurally protected or embedded foods than non-tool-assisted foraging techniques. This differs from necessity hypothesis in that it does not necessarily presuppose any differences in resource abundances between tool-using and non-tool using populations or species (Furuichi et al., 2015). Further, in contrast to the necessity hypothesis, the relative profitability hypothesis is primarily concerned with the maintenance of tool-using behaviors as a function of their increased economic utility, rather than their evolutionary origins.

The *limited invention hypothesis* is less commonly discussed, as it is difficult (or impossible) to empirically test (but see Furuichi et al. (2015)). It states that tool use may be invented rarely, or might be difficult to spread and/or maintain, and therefore the patterns of tool use we observe across species and populations are driven by neutral evolutionary processes rather than natural selection. This hypothesis predicts that instances of tool use might occur in disparate geographical areas where barriers to dispersal and cultural transmission exist, and that more frequent or complicated tool use will occur under conditions where constraints on transmission are relaxed. Barriers to dispersal and cultural transmission may be social, cognitive, or geographical. Fox et al. argued that the limited invention hypothesis is supported when no evidence for the first two can be found. However, it is possible that the origins of a behavior might be favored by selective pressures consistent with the necessity hypothesis or historical factors (the environment an animal evolved in) consistent with the opportunity hypotheses. Maintenance may be affected by factors consistent with the limited invention hypothesis.

Addressing these four hypotheses is challenging as they are not mutually exclusive. Within-species comparisons provide support for all of them (see Sanz and Morgan (2013) and Furuichi et al. (2015) for reviews). Within-population support in Brazilian populations of *Sapajus libidinosus* for the opportunity hypothesis has been found for nut cracking with stone tools at Fazenda Boa Vista (Spagnoletti et al., 2012) and using stones to dig for invertebrates and tubers at Serra da Capivara National Park (Falótico et al., 2017a). Between-species comparisons of chimpanzees and bonobos did not find support for the necessity, opportunity, or limited invention hypotheses and instead put forth that the ancestral ecological conditions in the Pleistocene might explain current behavioral variation (Furuichi et al., 2015). Current environmental conditions may not reflect the historic ecological or social conditions that favored the origins of tool use. The spatial scales of resource availability and territory size may differentially favor tool use in closely neighboring populations of the same species, so habitat-wide information may not be of use to particular social groups. The temporal scales in which resource bottlenecks occur that make tool-aided extractive foraging necessary for population persistence may also not be captured on the timescale of a typical research project. For these reasons it is important to be precise regarding if we are addressing the origins/and or maintenance of tool use, and endeavor to collect data on temporal or spatial scales that are most applicable to our hypotheses.

Another challenge about addressing these hypotheses is that the spatial and temporal scales at which we explore them may exist on different levels of biological organization (Mayr, 1961). Also, these hypotheses likely blur the lines between the proximate-ultimate distinction (Laland et al., 2011). The opportunity hypothesis, when viewed over evolutionary time is a question about the historic environmental conditions in which ancestral populations evolved and whether they were conducive towards tool use. However it is often treated as a hypothesis about proximate mechanisms: is stone tool use more likely to develop in social groups or species that are more likely to encounter stone tools in their environment? This contrasts with the *necessity hypothesis* and *relative profitability hypothesis* which focus on ultimate causation, ecological function, and fitness outcomes. The *limited invention hypothesis*, in contrast, focuses on the proximate factors which affect the spread of an adaptive tradition which may subsequently affect fitness outcomes. This interaction between both levels of biological organization is likely important for understanding the distribution of tool use behavior among primates.

### 1.3. Important factors in explaining variation in tool use in capuchins

Prior work has identified several factors which may explain variation in tool use among extant primates and what affects the origins and maintenance of this behavior (Fox et al., 1999; Fragaszy et al., 2004a; Moura and Lee, 2004; Sanz and Morgan, 2013; Visalberghi et al., 2005). These factors include resource limitation due to seasonal reductions in food abundance (i.e. tool use as a fallback strategy), high abundance of nutritious, embedded foods which require tools to access, low dietary richness, availability of stones and anvil sites, increased terrestriality and low predation risk.

Sufficient abundance of embedded foods and availability of tool-making materials are necessary preconditions for stone tool use. They provide a proximate explanation for why stone tool use is present (or absent) in a particular group or species. The abundance and nutritional quality of resources are also important in addressing the *relative profitability hypotheses*. Resource limitation, low dietary richness, increased terrestriality, and low predation risk impact the relative benefits and costs of tool use, providing an evolutionary explanation for why tool use arises and persists in some cases but not others.

The distribution of stone tool use among *Sapajus* populations supports the hypothesis that resource limitation and low dietary richness may favor the evolution of this behavioral strategy; tool using *Sapajus* populations tend to live in drier, more seasonal areas such as the Cerrado and Caatinga of Brazil which have less primary productivity, plant richness, and fruit abundance compared to populations living in the rainier, more productive, species rich Amazon forest (Moura and Lee, 2004). It has also been proposed to be a difference between gracile capuchins in central America, and robust capuchins in drier regions of Brazil. However, populations of *Cebus* live in diverse tropical ecosystems including seasonal dry forests (where the majority of well-studied populations live), primary and secondary rain forests, and montane cloud forests. To date, no studies have compared extractive foraging or tool use across these ecosystems within *Cebus*. The hypothesis that organisms will rely on tool use as a fallback strategy when easy to process fruits are seasonally limited (Marshall and Wrangham, 2007) receives support in robust capuchins and chimpanzees (Yamakoshi, 1998; Sanz and Morgan, 2013). However populations of *Cebus capucinus* rely more on extractive foraging of embedded insects and become more terrestrial when resources are limited at the end of the dry season (Perry and Ordoñez Jimenez, 2006; Melin et al., 2014).

The *relative efficiency hypothesis* has not been directly compared between populations of tool-using and non-tool-using capuchins. If a resource is abundant, easily encountered, and of particularly high quality, perhaps the costs of learning and using tools use will be compensated for by increased nutritional returns. Structurally protected foods which require tool use might be included in the diet if other resources in the environment are sufficiently rare or are of low nutritional quality in accordance with optimal foraging models (Emlen, 1966; MacArthur and Pianka, 1966). Additionally, resource abundance and quality is likely to interact with dietary richness, and an increased reliance on tool use might be favored when there are a limited set of food options available. However, any hypothesis about foraging optimality in primates might be better informed by social foraging models (Giraldeau and Caraco, 2000). Tool-use often generates producer-scrounger dynamics, which also affects the probability that a foraging behavior will be culturally transmitted (Ottoni et al., 2005). If age or sex differences in the efficiency or proclivity to use stone tools4exist, we might predict different reliance on tool use across age and sex classes and potential indirect impacts on social behavior.

Reduced dietary richness might also explain the difference between tool-using and non-tool-using populations of *Sapajus* and *Cebus*. Groups which live in areas with fewer potential diet items, such as islands or low quality habitats, may be forced to innovate and invest in potentially costly behavioral experimentation to make the best of a bad situation. It is also possible that differences in tool use rates might be influenced by ranging patterns. Gracile capuchins have larger home ranges than robust capuchins, on average (reviewed in Matthews (2009); Lynch Alfaro et al. (2014)), and thus might be able to access a more diverse set of food items that do not require tool-use compared to robust capuchins. However, it is also possible that groups of robust capuchins are just more capable of extracting resources out of a smaller area due to morphological specializations such as thicker enamel and larger molars (Lynch Alfaro et al., 2012).

Stones are needed for stone tools to be used. High availability of the necessary raw materials in the environment increases the probability that an individual will interact with them and innovate stone tool use. The importance of such opportunities for innovation is supported by within-*Sapajus* variation in stone tool use. Populations of *Sapajus* (and *Cebus*) living in the alluvial floodplains of the Amazon, for example, have less access to stones than populations living in rockier, drier habitats, and do not use stone tools (Moura and Lee, 2004). However, while the availability of raw materials may drive some within-genus or within-species variation, it is unlikely to explain broader taxonomic differences in the stone tool use as, within any *Cebus* or *Sapajus* species’ range, there exists considerable geological variation.

Increased terrestriality might explain the presence of stone tool use (Visalberghi et al., 2005; Meulman et al., 2012) consistent with the opportunity hypothesis. Individuals are more likely to encounter and wield stone tools on the ground than in the canopy. However, both *Cebus capucinus* and *Cebus albiforns* (Defler, 1979) can be exceptionally terrestrial, sometimes in response to seasonal fluctuations in fruit availability (Melin et al., 2014), yet, no previously studied populations of *Cebus* appear to use tools of any kind with any great frequency.

Decreased predation risk may lead arboreal species to become more terrestrial (Monteza-Moreno et al, in prep), thus, making them more likely to invent or use stone tools on the forest floor. However, the absence of predators might also affect ecological competition between conspecifics and heterospecifics with niche overlap. Increased within- and between-species competition may affect both population density and ranging behaviors of a candidate tool-using species and the abundance of heterospecifics with niche overlap, creating feedback looks which negatively affect species richness and abundance which may further favor the evolution of tool use. Reduced predation risk may also directly affect the probability of stone tool use by creating safe conditions where animals can afford exploratory behavior, potentially leading to the innovation of tool-use (van Schaik et al., 1999; Rutz et al., 2010). Stone tool use, in particular, may also directly attract predators; it is a loud, conspicuous activity that often occurs in predictable areas where resources are clustered, making tool-using individuals easy targets for predators. Stone tool use also requires concentration, thus decreasing an organism’s ability to be vigilant. Differences in predation risk may explain both within- and between-species variability in tool use.

## 2. STUDY SITE

Coiba National Park (CNP) in Gulf of Chiriqui of Panama is an archipelago encompassing 9 larger islands and more than 100 islets. The main island, Coiba, lies 23 km off Panama’s Pacific coast and has been geographically isolated from the mainland for 12-18,000 years (Ibáñez, 2011). CNP has significant ecological variation: moist and wet forest account for 60% of CNP (Castroviejo, 1997), and the park also includes mangroves and swamp forests. CNP receives 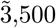 mm of rain per year which primarily falls during a marked wet season (May-December). CNP contains one of the largest remaining parcels of Pacific Central American forest; 75% of the park consists of mature forest, due to its historic (until 2004) role as a penal colony (Milton and Mittermeier, 1977). Coiba National Park is a UNESCO world heritage site and is a hotspot for marine and terrestrial plant and animal endemism (Ibáñez et al., 1997; Castroviejo, 1997; Olson, 1997; Zimmer, 1997; Duquette et al., 2017; Steinitz et al., 2005), much of which is understudied. This includes five endemic species and subspecies of terrestrial mammals: the Coiban agouti *Dasyprocta coibae*, Coiba Island or Rothschilds’ white-tail deer *Odocoileus virginianus rothschildi*, Coiba island howler monkey*5Alouatta palliata coibiensis* or *Alouatta coibensis*, black-eared opossum *Didelphis marsupialis battyi*, and potentially a rodent of the genus *Zygodontomys* (González et al., 2010).

Three of the largest islands in CNP-Coiba (50,314 ha), Jicarón (2,002 ha), and Ranchería (222 ha) support populations of capuchin monkeys (*Cebus capucinus imitator*) (Figure S1). Capuchins living on the islands within the Coiban archipelago provide a natural laboratory for examining hypotheses about the evolutionary drivers of tool use. Populations are geographically isolated by the ocean, strongly limiting genetic flow and cultural transmission. Additionally, the three islands differ in the number of potential food items available to capuchins, following well-established patterns of species richness varying as a function of distance from the mainland and island area (MacArthur and Wilson, 1967). Transect data (Ibáñez et al, in prep) shows that Coiba, the largest island, located in the middle of the archipelago, is home to 1001 plant species. Jicarón (the most distant island) and Ranchería, in contrast, host similarly depauperate plant communities (261 and 262 species, respectively), despite a nearly 10-fold difference in size (Ibáñez, 2001). Smaller islands are also more likely to suffer from ecological perturbations due to edge effects. These two factors combined with the highly terrestrial behavior of capuchins in CNP (which may increase encounter rates with material for stone tool use) due to an absence of terrestrial predators on the islands (Milton and Mittermeier, 1977) provides ample opportunity (and decreases potential costs) for capuchins to use stone tools.

## 3. METHODS

### 3.1. Coastal surveys

In 2004, A. Ibáñez first observed capuchins using stone tools to crack open the endocarps of *Terminalia catappa*− a coastal tree from Asia which has been nativized in the Neotropics and is known commonly in Spanish as *almendro de playa* and in English as *sea almonds*. Following up on this initial report, a documentary field crew captured this nut cracking behavior on film in 2013 (Paul Stewart and Huw Cordey, personal communication). To further document and determine the extent of this unique behavioral variant, we conducted field surveys from March 2017-March 2018. B. Barrett and C. Monteza surveyed the coasts of the three islands in CNP with capuchin populations by boat to identify *T. catappa* groves. We then surveyed these areas (including the entirety of Jicarón and Ranchería) on foot to look for evidence of tool use, including broken endocarps, hammerstones and anvils, cracked crab and bivalve shells, and coconuts.

### 3.2. Camera Trapping Effort

To assess potential tool use sites, we deployed unbaited photo (Hyperfire HC600) and video (Ultrafire XR6) camera-traps (Reconyx, Inc, WI, USA) between March 25, 2017 and March 26, 2018. Photo-trapping is advantageous for recording localized tool use for unhabituated primates in remote areas (Sanz et al., 2004). Photo camera-traps were set to take a series of 10 images per trigger event with no delay between triggers, allowing us to document behavioral sequences. Video camera traps captured sequences consisting of one photo followed by 30 seconds film per trigger. We deployed cameras at 35 sites on the islands of Coiba (4 sites, 532 trap-nights), Jicarón (30 sites, 3466 trap-nights), and Ranchería (1 site, 148 trap-nights). At four sites (2 Coiba, 1 Ranchería, 1 Jicarón) where we found almendros and capuchins but few rocks, we set up experimental anvils and hammerstones to see if capuchins would use them.

### 3.3. Weighing and scanning tools

Graphic documentation in the field consisted of geo-located digital pictures of sites and tools, 3D models of sites using geo-referenced images (Agisoft Photoscan), hand-made and digital mapping, and situational sketches. The use of this set of standardized archaeological methods facilitates comparison with archaeological sites (and any others when the same protocols are utilized). Using a digital scale, we weighed a subset of the hammerstones (N=60) used in the behavioral sequences captured by our camera traps or observed during coastal surveys (i.e. stones which were pile on broken *Terminalia* endocraps, resting on coastal driftwood). Broken hammers were refitted on site before being weighted. A set of photographs of several of these tools were taken with a goal to create 3D models according to protocols established by Porter et al. (2016).

### 3.4. Statistics

To estimate the size of stone tools and monthly rates of tool use across camera stations we used generalized linear models and generalized linear mixed effect models fit using the map2stan function in the rethinking package (v. 1.59) (McElreath, 2016) in R (v. 3.43). map2stan is a front-end which utilizes rSTAN (v. 2.17.2), a Hamiltonian MCMC sampling engine (Stan Development Team, 2016). To estimate mean stone tool size, we fit a Gamma GLM using a log link as stone tool weights are positive, non-normal, and lower-bound by zero. To estimate monthly tool use rates we used an aggregated binomial GLMM with a logit link. To analyze stone tool use rates we analyzed a subset of camera trap data where stone tool use was observed at least once. Our outcome was the number of days per month where tool use was observed at each camera station and the number of trials was the number of days per month where a camera recorded a capuchin at least once. We estimated varying intercepts for each camera station (N=7), camera station deployment (N=13), and unique month (N=12). All data, model, and graphing code for this paper can be found at this manuscript’s corresponding GitHub repository. These annual rates are a coarse, preliminary measure of stone tool use in this population and future analyses will account for inter-individual and monthly variation in tool use rate at more consistently sampled tool use sites.

## 4. FINDINGS

### 4.1. Habitual reliance on stone-tool aided extractive foraging in white-faced capuchins

We found that capuchins on Jicarón in CNP habitually use stone tools to open a variety of food resources including *Terminalia catappa* endocarps (Figure 2, Supplemental Video), hermit crabs (Figure 4c, Supplemental Video), Halloween crabs (*Gecarcinus quadratus*) (Figure 4b,d), coconuts (*Cocos nucifera*) (Figure S4), and marine snails. Middens of *T. catappa* shells, coconut husks, and tool shards are readily apparent at tool use sites (Figure 4b, S3c), although coconuts are almost always opened using only an anvil, as has been described on Coiba Island (Méndez-Carvajal and Valdés-Díaz, 2017). Elusive tool use occupations are also commonly observed in the intertidal and are regularly wiped-out by daily changes in the tide (Figure S3a). Small and medium sized occupations are located within dry stream beds (Figure S3b) and larger accumulation of processed food lie on higher ground (Figure S3c). This situation suggests that the joint effect of the tide and seasonal activity of the stream have an impact on site formation processes and thereby, may affect site visibility. Further, during coastal surveys, we observed stones piled on broken clam shells at the edge of the intertidal zone, consistent with capuchins using stone tools to exploit this rich marine resource (Figure S2). In some instances, the stones were spaced in a regular pattern, at distances similar to the inter-individual distances observed in foraging capuchins. Destruction of cameras by vandals and harsh marine conditions prevented us from confirming this behavior.

**FIGURE 1.**
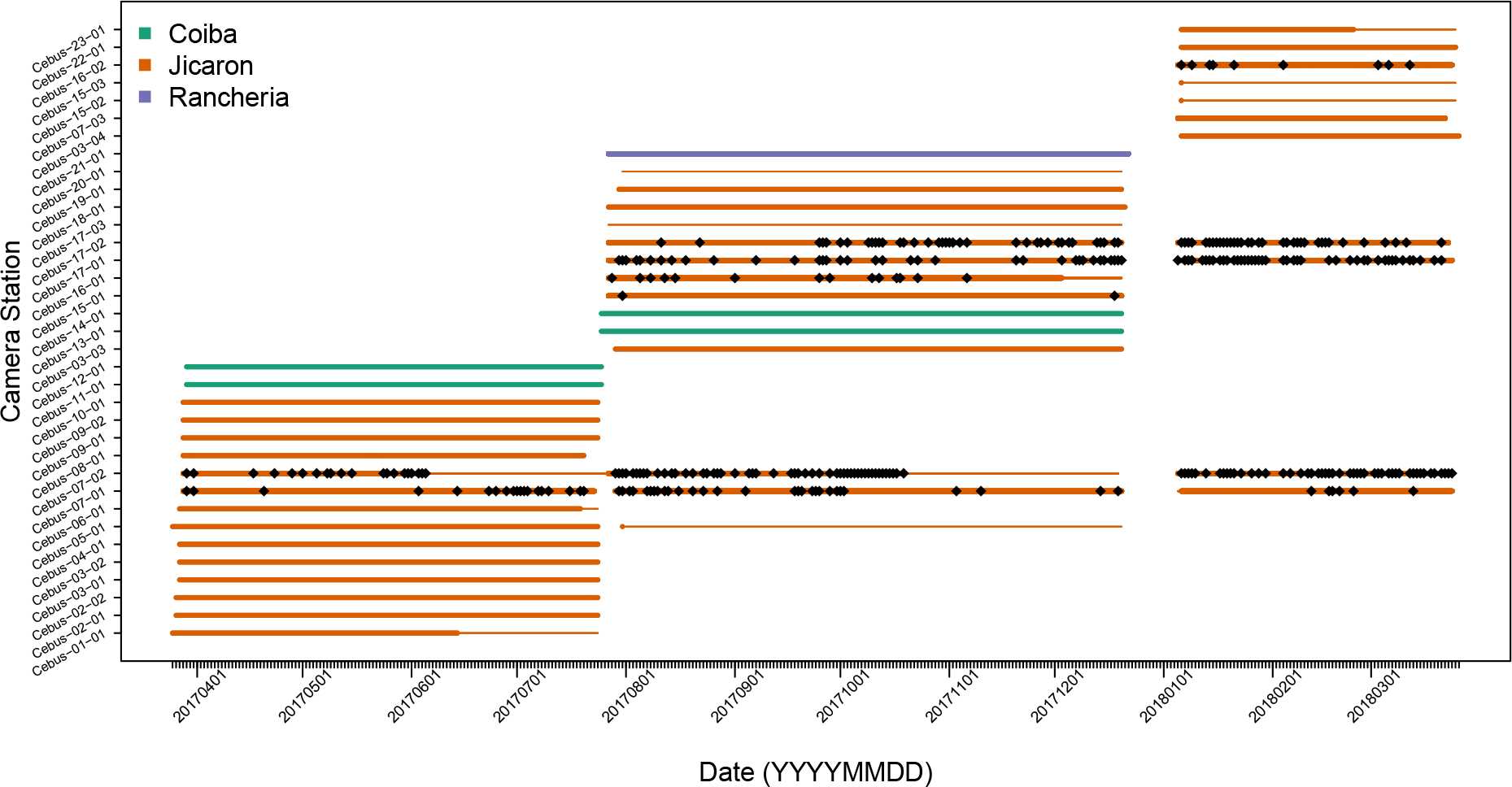
Visualization of camera trap sampling effort. Rows are camera trap locations. Smaller lines indicate length of deployment, thick lines indicate period of deployment when camera was filming and focused on tool use site (i.e was not moved by animals, had working batteries, was not stolen or vandalized). Diamond points occur at each day where stone tool use was recorded. Colors indicate island in CNP.

**Figure 2.**
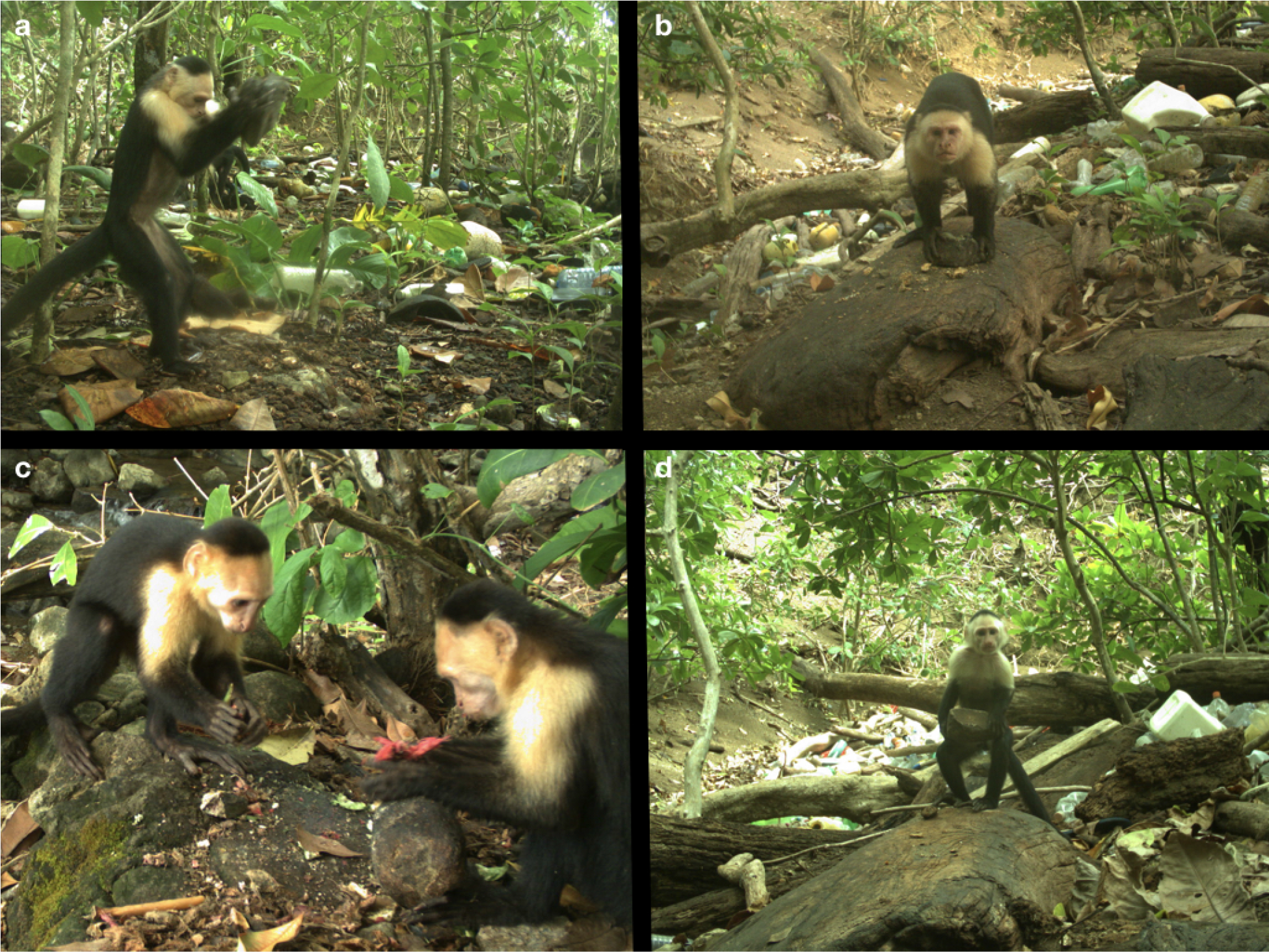
A juvenile male capuchin uses a hammerstone to crack open a *T. catappa* en-docarp on a stone anvil (a). An adult male after cracking open a *T. catappa* endocarp on a wooden anvil. Juvenile male capuchin observing an older juvenile processing *T. cat-appa* endocarps with a hammerstone (c). Juvenile male about to process *T. catappa*- note prehensile tail used for support.

In contrast to Jicarón, our initial coastal surveys and camera trapping failed to return evidence of stone tool use by capuchins on the islands of Coiba or Ranchería, despite the presence of the necessary raw material ( *Terminalia catappa* and stones) at both sites. We did not observe tool use at any of the artificial anvil sites we set up on the islands (Figure 1).

### 4.2. Description of stone tools

Capuchin tool use primarily consisted of pounding food items using a hammerstone. Preliminary observations indicate that the pounding technique is direct percussion with hard hammer (*sensu* Inizan et al. (1999), Figure 2, Supplemental Video). Food items were placed on stationary anvils of mineral or organic material (Figure 4). Mineral anvils included rocky outcroppings and bedrock along stream banks, in the forest interior and in the intertidal zone (Figure S3). In cases where stones were used as anvils, this technique could have produced a rebound force. Large fallen trees and logs in the forest and washed up in the intertidal zone were also used for pounding, as were the bases of trees and tree branches. In some cases, hammerstones were transported 2-3 m up into the canopy. When used as a support for pounding, wood is less likely to produce a rebound force, suggesting a use of wood as a stand-on surface to increase the accuracy of the strike or easy collection of the food item.

**FIGURE 3.**
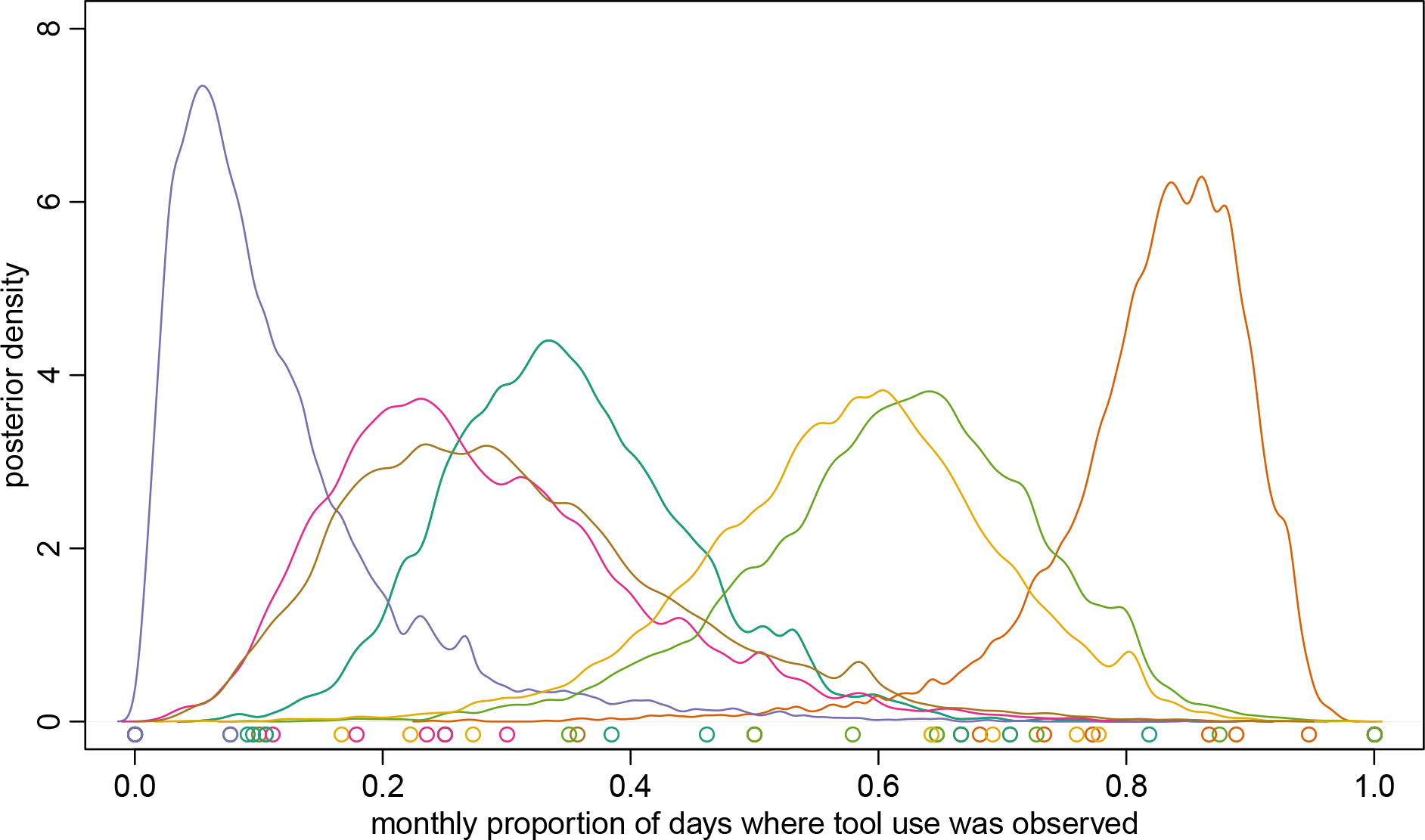
Tool use sites differed greatly in their observed usage. On average, tool use was observed at each camera site on 45.3% of days where capuchins were observed. Cameras at the most active tool use sites (orange curve on far right) recorded tool use on 83.2% of the days where capuchins were observed. Curves show posterior predictions of probability of observing tool use on each day conditional on observing a capuchin across all months. Colors correspond to camera stations. Points on X-axis are raw proportions.

**FIGURE 4.**
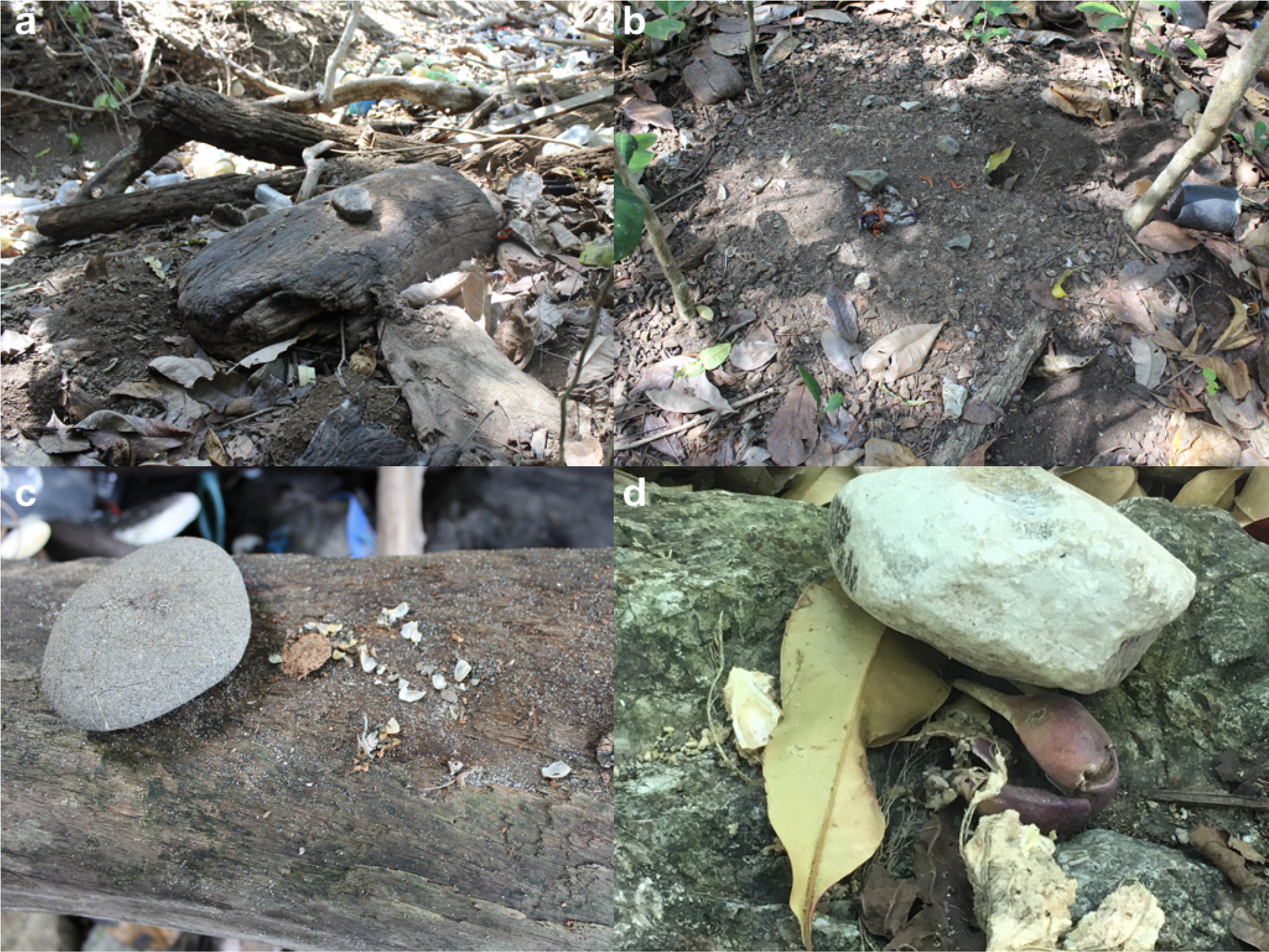
Wooden (a,c) and stone anvils (b,d) used by capuchins to process structurally protected foods. Note accumulation of processed materials including old endocarps, coconut husks, and shells in a and b. Panel c shoes hermit crab exoskeleton and shell remains processed with a stone tool on dead tree branch. Panels b and d show stone hammer on stone anvil with *T. catappa endocarps* and Halloween crab remains. The leaf stuck under the hammer in panel d indicates that the food was processed shortly before discovery. Percussion impacts can be observed on the crab limb.

Capuchins primarily used cobbles collected from streams as hammerstones. In some cases, stones from the marine intertidal and eroded hillsides were also used to process food items. A sample of the tools randomly collected on trips in July 2017 and January 2018 from 60 tool use sites used by capuchins on Jicarón varied greatly in mass (mean=740.7 g, sd=402.7, range=172-2256 g, N=60; Figure 5). Capuchins used large stones-one weighing > 2 kg-to process *Terminalia catappa*, adopting a two-handed grip with a bipedal stance, and often using their semi-prehensile tails for support and leverage. In contrast, smaller items such as hermit crabs and snails were cracked with a two or one-handed grip in a crouching position.

**FIGURE 5.**
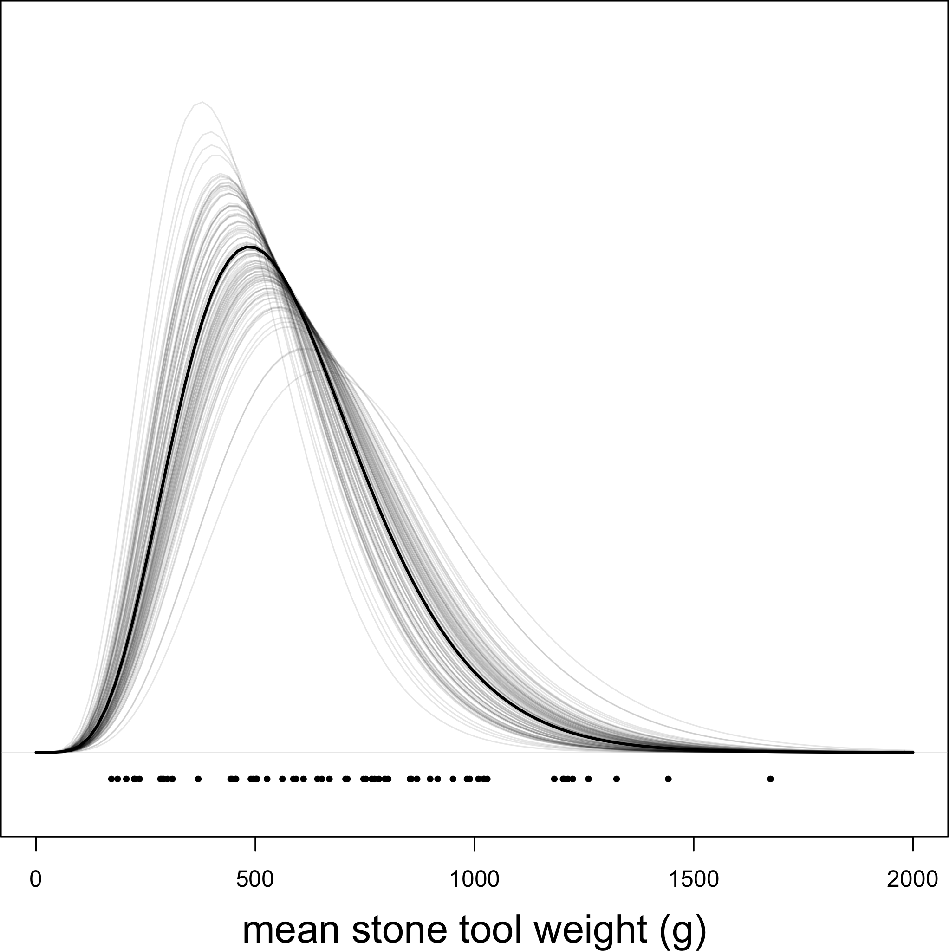
Model predictions from a gamma GLM of mean stone tool weight. Dark line is posterior mean estimate, while lighter lines are 100 randomly sampled posterior predictions to visualize uncertainty.

Based on our photographic evidence and surveys, both hammerstones (Figure S5) and food items were sometimes transported to the tool use sites (Supplemental Video). *T. catappa* endocarps, for example, were often collected multiply underneath *T. catappa* trees at the shoreline and transported to anvil sites deeper in the forest. In this population, the use of stone tools is heavily male-biased. Over the course of one year we did not observe a single adult female use stone tools despite them commonly engaging in scrounging behavior in the vicinity. Tool use in this population also generates interest and close-range observation from conspecifics, providing opportunities for social learning (Figure 2c).

## 5. DISCUSSION

This is the first description of habitual stone tool use in gracile capuchin monkeys, adding *Cebus* to the list of primate genera with populations who habitually use stone tools in the wild (*Sapajus, Macaca, Pan*). Tool use by capuchins on Jicarón shares key features with the patterns of behavior described in longtailed macaque populations in Thailand: stone tool use occurs solely on islands and is focused on coastal resources (Gumert and Malaivijitnond, 2012). It is also similar to chimpanzees in that capuchins transport both processing material and stone tools to particular anvil sites that are reused over time. However, tool use also appears more sporadically in space. This population of *Cebus*, several populations of long-tailed macaques in Thailand (Gumert and Malaivijitnond, 2012; Falótico et al., 2017b), and some populations of *Sapajus* (Aguiar et al., 2014) target one of the same species- *Terminalia catappa*. This provides a unique opportunity to directly compare tool use behavior, quantify differences in foraging efficiency, material choice, and the social behaviors surrounding tool use for an identical resource across three primate species in the wild.

Stone tool use appear to have some potential seasonality-the lowest rates of tool use occurred in the transition periods between wet and dry season (Figure 7). This pattern differs from what has been observed with extractive foraging behaviors in *Cebus* in neotropical dry forests (Perry and Ordoñez Jimenez, 2006; Melin et al., 2014) where extractive foraging increases in the transition periods between seasons. More rigorous analysis of both behavioral and ecological data in a comparable manner is needed to verify this observation.

**FIGURE 6.**
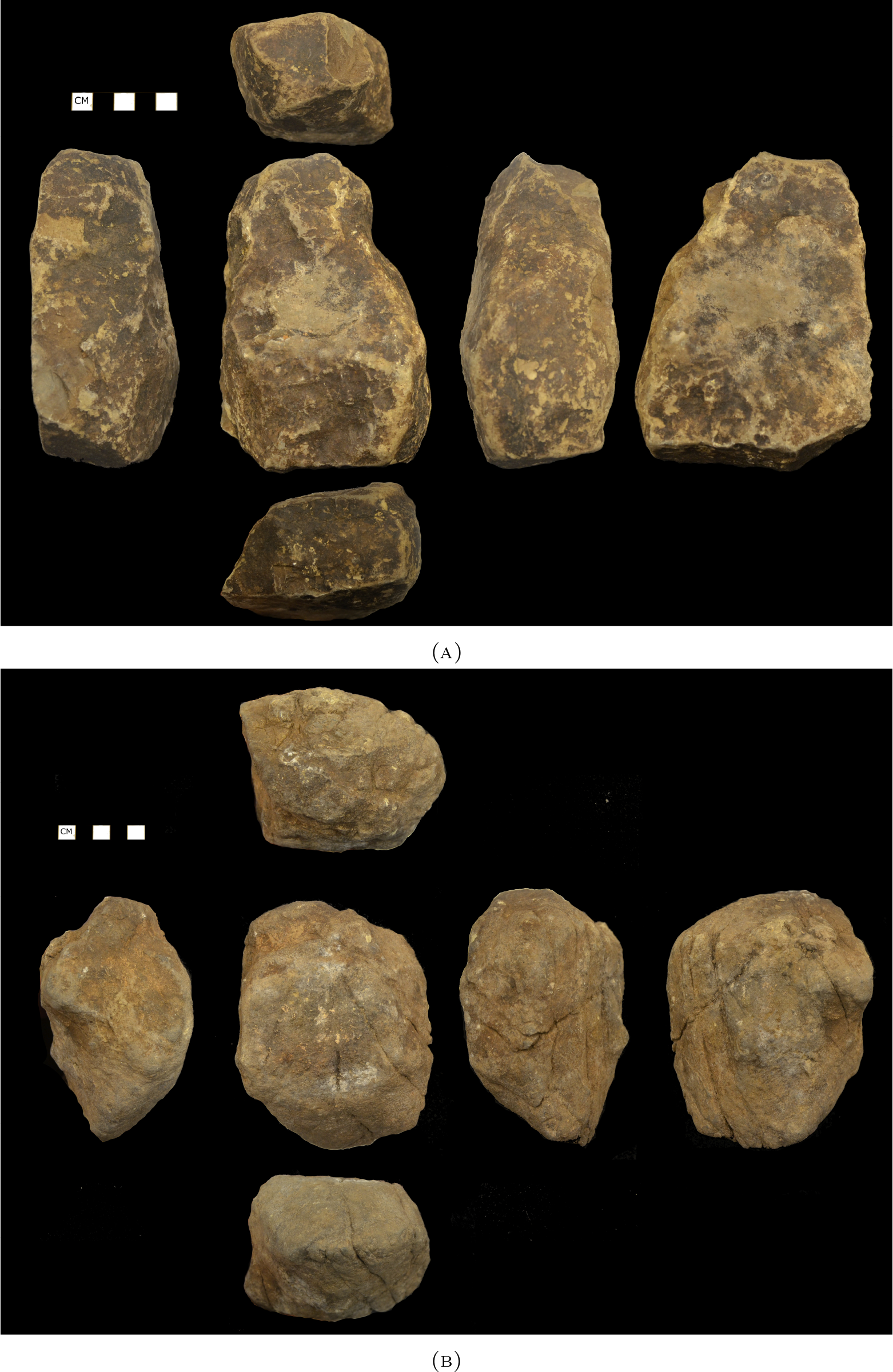
Examples of hammerstones from Jicarón sites used for nut cracking weighing (a) 989 g and (b) 1202 g.

**FIGURE 7.**
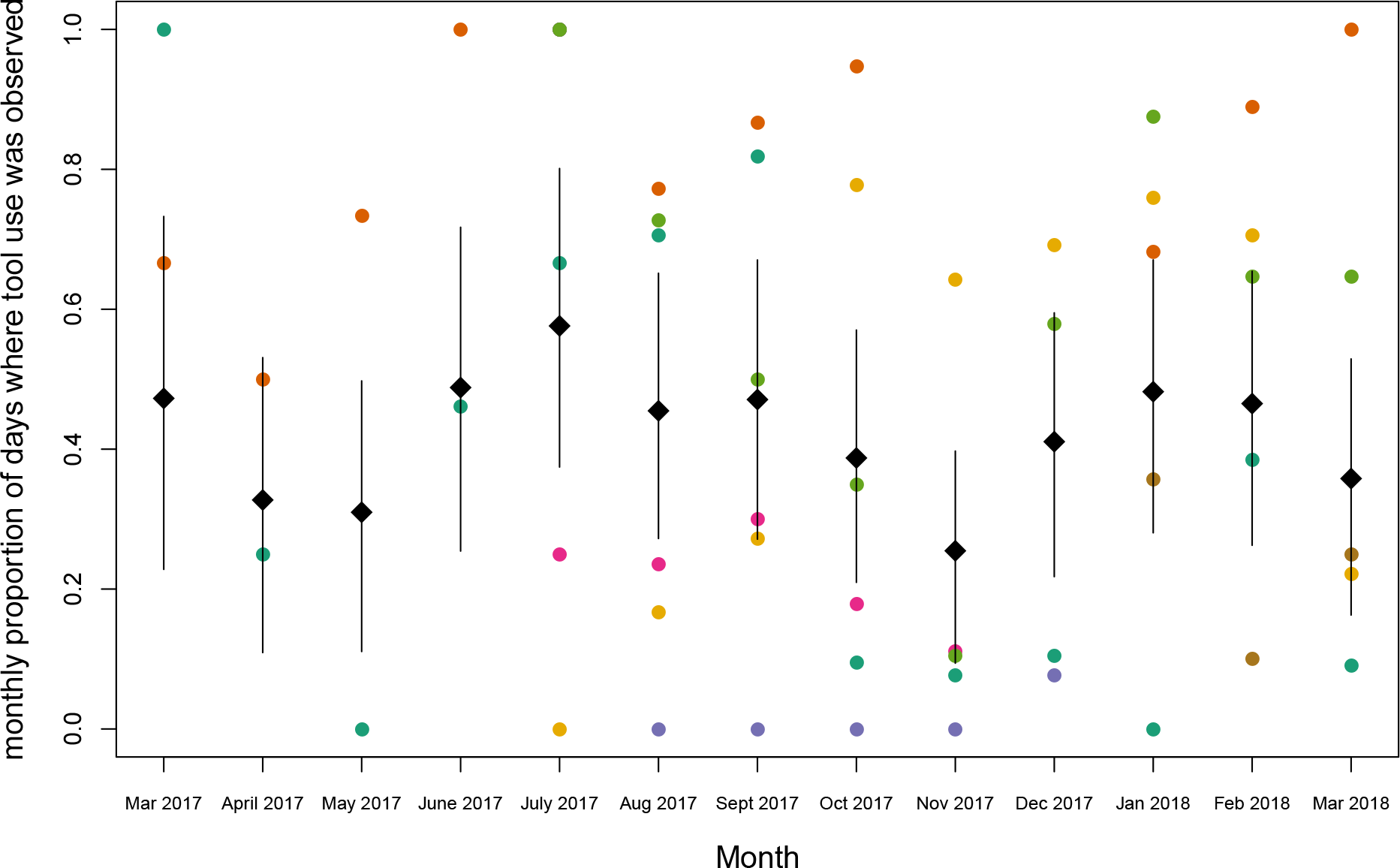
Tool use rates appear lowest in the transition months from dry to wet (April and May) and wet to dry (Nov) seasons. Graph shows model predictions of the probability of observing tool use per day for each month conditional on observing a capuchin at each camera site. Gray diamonds lie at posterior mean estimate while lines indicate 89% posterior credible interval. Each color is a raw proportion per camera deployment which corresponds with a camera trap station.

The patterns of tool use observed in capuchins on Jicarón also share many similarities with nut-cracking behavior in *Sapajus*. Individuals habitually reused tools and favored a small number of repeatedly visited tool use sites to which they transported tools and materials for processing. The accumulation of debris at these tool use sites and the comparatively dry climate on Jicarón offers potential for preservation and archaeological excavation.

### 5.1. Ecology of stone tool use

Capuchins in CNP spend an unusually high percentage of their time traveling and foraging on the ground (Milton and Mittermeier, 1977), which may be due to the absence of terrestrial predators on these oceanic islands (Monteza-Moreno et al., in prep). By providing opportunities for capuchins to interact with and manipulate stones, this expansion into the terrestrial niche may have facilitated the emergence of this plausible cultural tradition. Additionally, the islands of CNP are particularly prone to climatic variation from ENSO cycles (Wang and Fiedler, 2006), edge effects, and climate change. These factors, combined with a decrease in dietary richness might drive foraging innovation in times of resource scarcity. However, it is important to note that stone tool use on Jicarón appears to be limited to particular sections of the island and likely to a single group, despite the availability of the necessary resources in other areas. Although surveys of the coastline and forest interior of Coiba are still underway, we have yet to find any evidence of stone tool use on this island or on Ranchería. As noted above, site preservation might have an impact on the visibility of tool-use but it is unlikely to obliterate all signs of such behavior. Further, many of the resources which capuchins on Jicarón process using stone tools are available to and 3aten by other white-faced capuchin populations including hermit crabs (Soley et al., 2017), Halloween crabs (B. Barrett, personal observation), marine snails and coconuts (Méndez-Carvajal and Valdés-Díaz, 2017). Fleshy exocarps of *T. catappa* are consumed by other capuchin groups across Coiba National Park and in Costa Rica (Campbell, 2013)− however capuchins are unable to access the nutritious endocarps without tools. This begs the question: why does this variation exist? How does between-group and between-island dispersal structure cultural and genetic variation?

### 5.2. Conservation implications

How isolated populations fare is a major conservation concern, and island systems provide an opportunity to explore the impacts of long-term isolation (Castroviejo, 1997) and small population size while providing insight into how rapid anthropogenic change may affect threatened mammal populations due to species invasions (Schofield, 1989) and climate change (Bergstrom and Chown, 1999; Courchamp et al., 2014). The capuchins of Coiba National Park are an excellent study system to understand how geographical isolation affects both genetic and cultural diversity, and the ability of these two interacting inheritance systems to facilitate species' responses to environmental change. Evidence of amelanism for capuchins on the island of Coiba (Duquette et al., 2017) are consistent with increased homozygosity due to founder effects or inbreeding. Yet the observed behavioral flexibility of capuchins on Jicarón displayed in this paper and of capuchins on Coiba (Méndez-Carvajal and Valdés-Díaz, 2017) suggest that they are behaviorally adapting to island life-potentially through social learning. Continued investigations into the behavioral flexibility of CNP capuchins through foraging innovation or behavioral and social flexibility will provide insight into the contributions of genetic and cultural variability in responding to ecological stress in a primate population.

### 5.3. Taxonomic status

Lastly, evaluating the taxonomic status of capuchin monkeys living in Coiba National Park is essential. Knowing whether these capuchins are an endemic species or subspecies will be important in affecting conservation decisions in an area that is threatened by development and climate change (Steinitz et al., 2005). Coiba is home to an endemic subspecies of howler monkey, *Alouatta palliata coibensis* (Cortés-Ortiz et al., 2015; Ellsworth and Hoelzer, 2006), which some consider to be its own species, *Alouatta coibensis* (Thomas, 1902; Méndez-Carvajal, 2012). However, no similar genetic or morphological comparisons of CNP versus mainland capuchin populations have been undertaken to determine if the Coiban capuchins are taxonomically unique. Capuchins samples collected in Coiba National Park in the early 20th century were smaller than mainland samples (see (Goldman, 1920); pp. 232-233), an observation consistent with the “island rule”- that larger mammals tend to become smaller on islands to cope with resource limitation (Foster, 1964; Lomolino, 1985). Analysis of the genetic diversity of capuchins living across the archipelago and those of the nearby Panamanian mainland will help address taxonomic status and permit us to understand if social systems flexibly respond to the ecological pressures associated with island living. Conserving behavioral variation of cultural traditions in capuchins is also important to inform conservation decisions in CNP, as culture is an important means by which organisms adapt to environmental change (Ryan, 2006; Whitehead, 2010). Panama is also an important biogeographical corridor in the evolution of *Cebus*, and genetic analysis of capuchins across Panama will clarify whether Pacific capuchins should be considered a separate species (*Cebus imitator*) from their Atlantic congeners (*Cebus capucinus*) (Lynch Alfaro et al., 2012, 2014).

## ACKNOWLEDGEMENTS

This research was funded by a Coss Award for International Field Research, a Smithsonian Tropical Research Institute Short-term Fellowship, and L.S.B. Leakey Foundation grant awarded to BJB as well as funds from the Max Planck Institute. It was also supported by a Packard Foundation Fellowship (2016-65130) and grant from the National Science Foundation (NSF BCS 1514174) awarded to MCC. Thanks to Owen McMillan and the staff at STRI’s Ranchería field station for support; Eliezer Vega, Chris Dillis, Pedro Luis Castillo, Zarluis Miguel Mijango, and Evelyn del Rosario assisted with field work. Thanks to Paul Stewart and Huw Cordey of Silverback films. This research was conducted under Ministerio de Ambiente, Panamá, scientific permit No. SC/A-23-17 and corresponding renewals and addenda.

---

**Figure S1.**
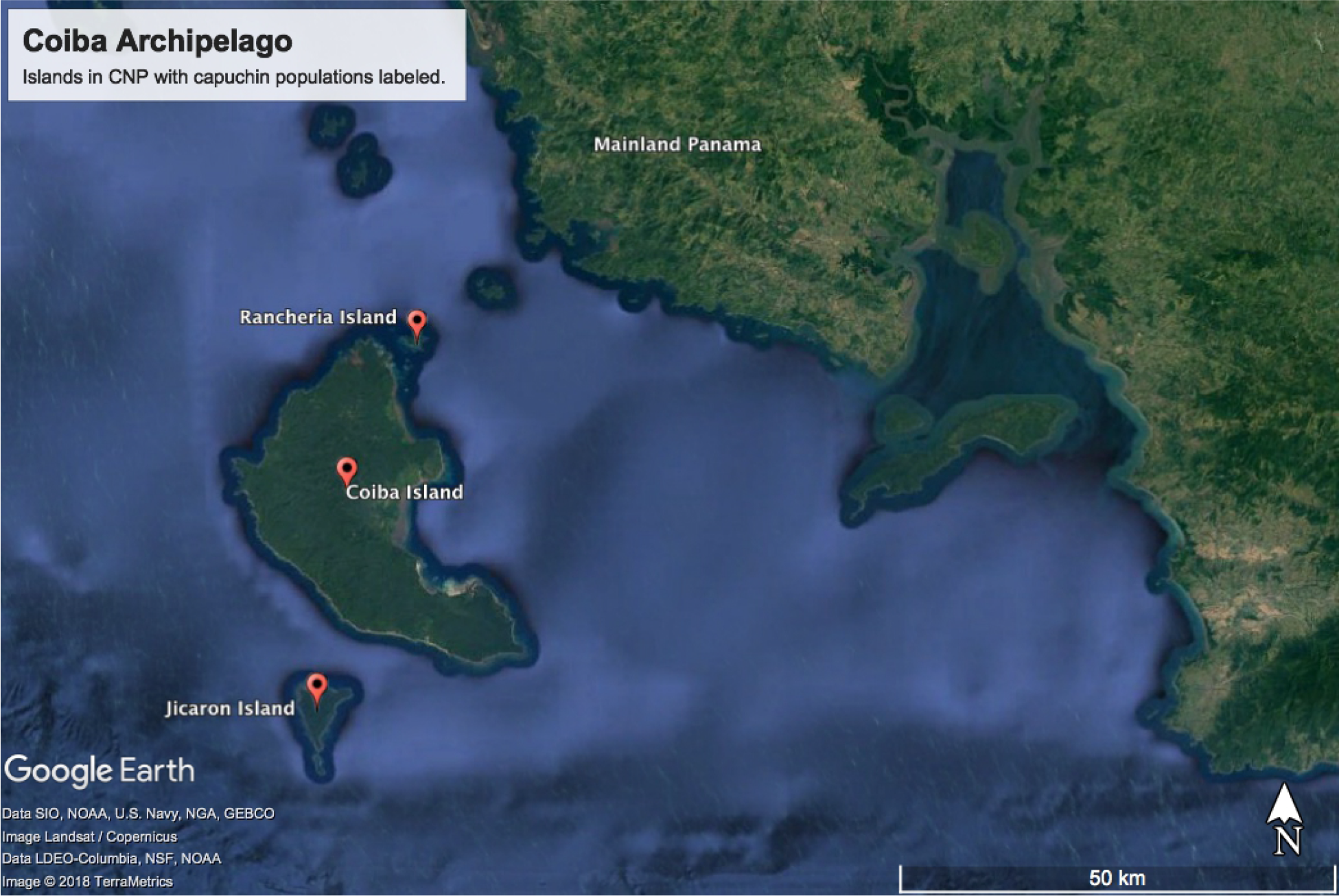
Map of the Coiba Archipelago off the Pacific coast of Panama. The three islands hosting populations of capuchin monkeys in CNP are labeled.

**Figure S2.**
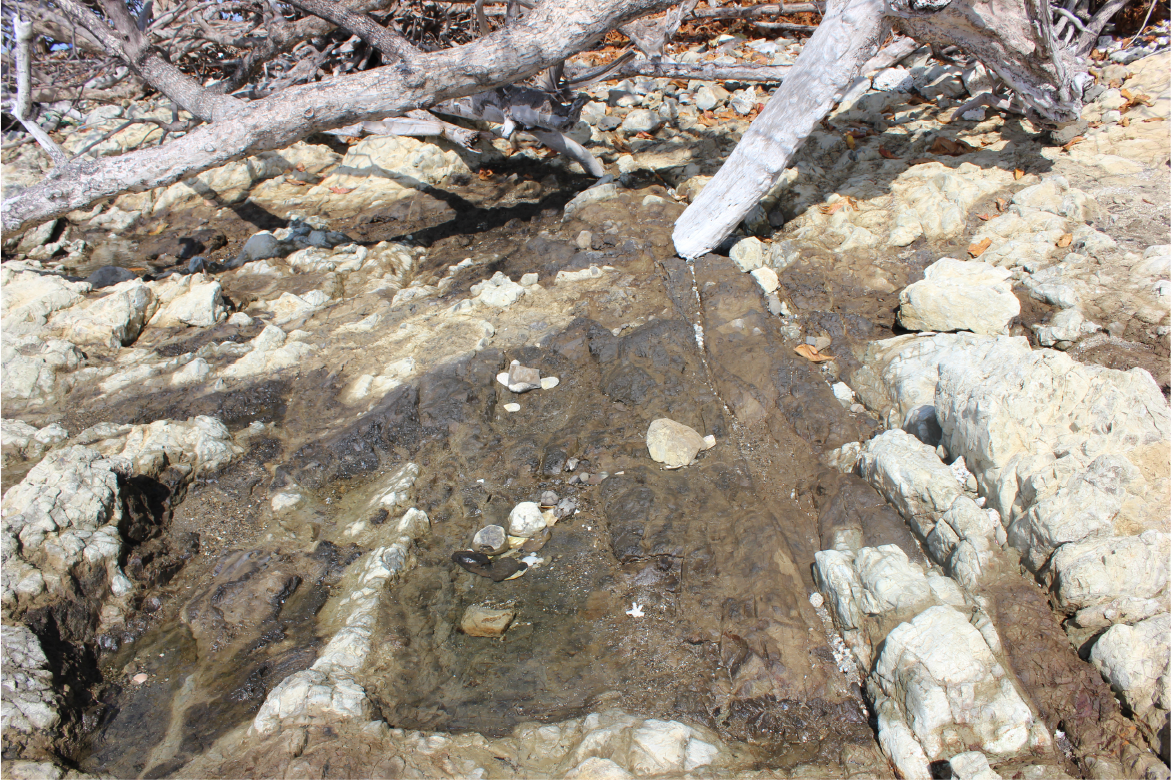
Recently broken clamshells at edge of intertidal zone with stones piled on them. Stone and clam accumulations spaced at distances consistent with between individual capuchin foraging proximities. We are unsure if capuchins are responsible for this or if this is a natural accumulation from tidal processes.

**Figure S3.**
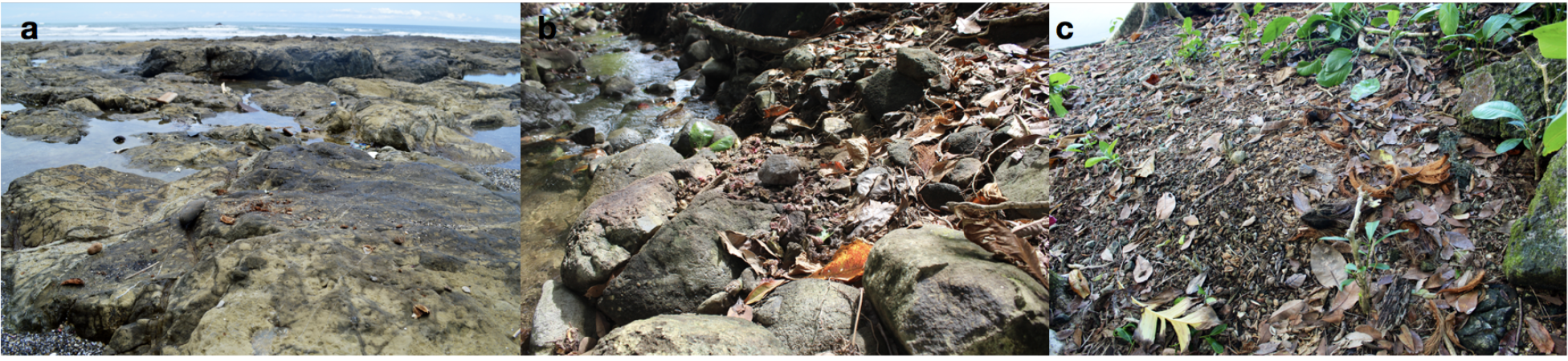
(a) Elusive tool use occupation in the intertidal zone that is regularly destroyed by daily tidal changes. (b) Medium sized tool use occupation along stream bank that is destroyed less regularly by seasonal floods. (c) Large tool use occupation lying on higher ground away from streams and shore. Accumulation of tools and processed food shows potential for archaeological excavation.

**Figure S4.**
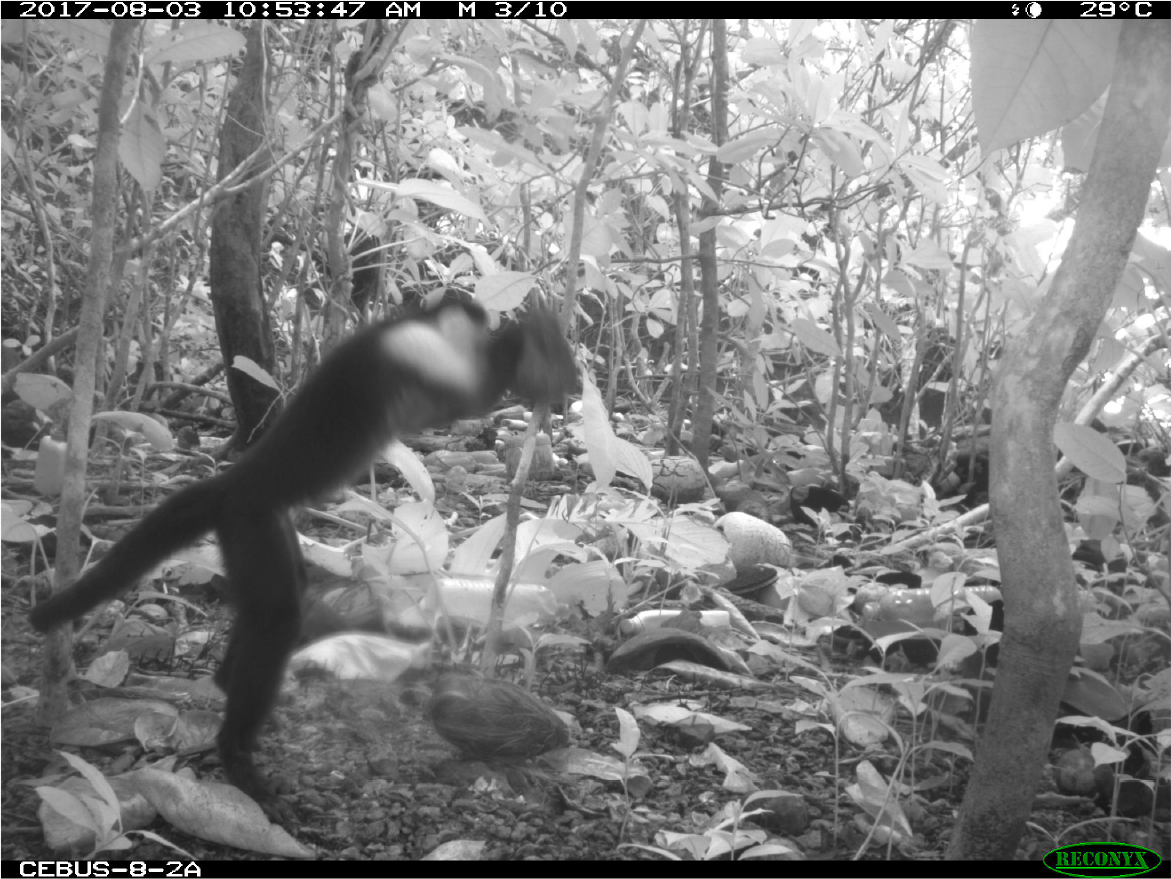
A juvenile male capuchin monkey breaking open a coconut husk with a hammerstone on a stone anvil.

**Figure S5.**
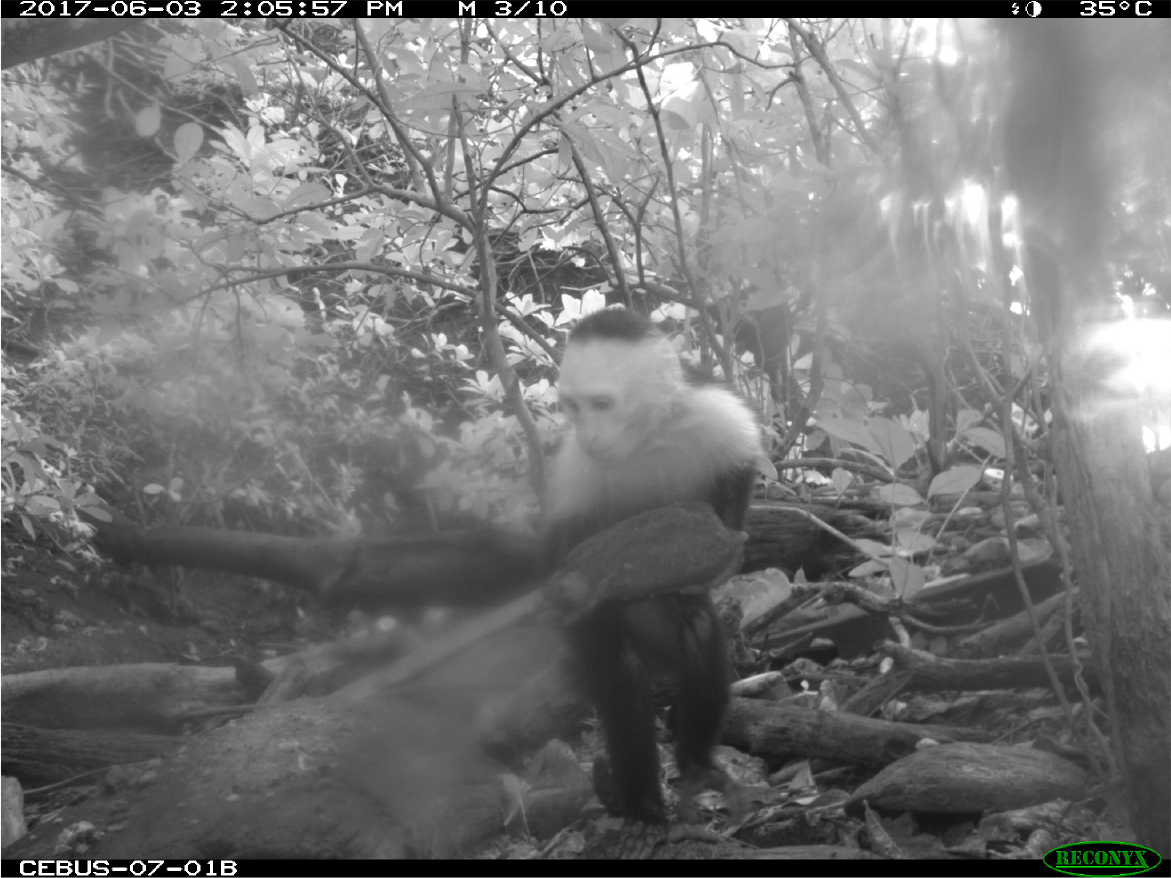
A juvenile male capuchin monkey transporting a stream cobble to a wooden
anvil on the forest edge.

**Table S1.** Model parameter predictions for monthly tool use rates. Varying effects are offset form overall intercept, *a*.

| Parameter | Posterior Mean | Posterior SD | 89% PCI |  |
| --- | --- | --- | --- | --- |
| $a_{Mar2017}$ | 0.05 | 0.59 | -0.86 | 0.97 |
| $a_{April2017}$ | -0.64 | 0.56 | -1.51 | 0.23 |
| $a_{May2017}$ | -0.72 | 0.51 | -1.50 | 0.11 |
| $a_{Jun2017}$ | 0.13 | 0.50 | -0.64 | 0.94 |
| $a_{Jul2017}$ | 0.51 | 0.45 | -0.18 | 1.25 |
| $a_{Aug2017}$ | -0.03 | 0.37 | -0.59 | 0.57 |
| $a_{Sep2017}$ | 0.04 | 0.38 | -0.51 | 0.69 |
| $a_{Oct2017}$ | -0.32 | 0.36 | -0.83 | 0.29 |
| $a_{Nov2017}$ | -0.98 | 0.41 | -1.61 | -0.32 |
| $a_{Dec2017}$ | -0.23 | 0.39 | -0.86 | 0.35 |
| $a_{Jan2018}$ | 0.09 | 0.39 | -0.50 | 0.69 |
| $a_{Feb2018}$ | 0.02 | 0.40 | -0.58 | 0.65 |
| $a_{Mar2018}$ | -0.45 | 0.39 | -1.07 | 0.19 |
| $a_{Cebus-07-01}$ | -0.45 | 0.60 | -1.38 | 0.46 |
| $a_{Cebus-07-02}$ | 1.79 | 0.68 | 0.76 | 2.86 |
| $a_{Cebus-15-01}$ | -2.12 | 0.96 | -3.65 | -0.58 |
| $a_{Cebus-16-01}$ | -0.82 | 0.73 | -2.00 | 0.28 |
| $a_{Cebus-17-01}$ | 0.69 | 0.64 | -0.42 | 1.61 |
| $a_{Cebus-17-02}$ | 0.52 | 0.64 | -0.48 | 1.46 |
| $a_{Cebus-16-02}$ | -0.70 | 0.76 | -1.89 | 0.46 |
| $a_{Cebus-07-01A}$ | 0.05 | 0.50 | -0.72 | 0.85 |
| $a_{Cebus-07-01B}$ | -0.10 | 0.47 | -0.90 | 0.59 |
| $a_{Cebus-07-01C}$ | -0.63 | 0.53 | -1.43 | 0.20 |
| $a_{Cebus-07-02A}$ | -0.22 | 0.54 | -1.09 | 0.58 |
| $a_{Cebus-07-02B}$ | 0.07 | 0.56 | -0.76 | 0.99 |
| $a_{Cebus-07-02C}$ | 0.00 | 0.55 | -0.83 | 0.91 |
| $a_{Cebus-15-01B}$ | -0.57 | 0.70 | -1.63 | 0.48 |
| $a_{Cebus-16-01B}$ | -0.34 | 0.58 | -1.26 | 0.54 |
| $a_{Cebus-16-02C}$ | -0.31 | 0.57 | -1.22 | 0.58 |
| $a_{Cebus-17-01B}$ | -0.53 | 0.51 | -1.32 | 0.27 |
| $a_{Cebus-17-01C}$ | 0.29 | 0.56 | -0.58 | 1.16 |
| $a_{Cebus-17-02B}$ | -0.15 | 0.51 | -0.90 | 0.67 |
| $a_{Cebus-17-02C}$ | -0.10 | 0.51 | -0.93 | 0.70 |
| $a$ | -0.19 | 0.25 | -0.56 | 0.19 |
| $\sigma_{month}$ | 0.59 | 0.22 | 0.24 | 0.89 |
| $\sigma_{camstat}$ | 1.60 | 0.66 | 0.67 | 2.53 |
| $\sigma_{camstatdeploy}$ | 0.56 | 0.31 | 0.07 | 0.93 |

**Table S2.** Model parameter predictions of Gamma GLM with a log-link.

| Parameter | Posterior Mean | Posterior SD | 89% PCI |  |
| --- | --- | --- | --- | --- |
| $a$ | 6.51 | 0.05 | 6.44 | 6.58 |
| $scale$ | 88.29 | 8.36 | 74.43 | 100.69 |

